# Parameterization of regulatory nodes for engineering broad host range heterologous gene expression

**DOI:** 10.1101/2021.05.27.445938

**Authors:** Pablo I. Nikel, Ilaria Benedetti, Victor de Lorenzo, Belen Calles

**Author notes:** Correspondence to: Víctor de Lorenzo, Centro Nacional de Biotecnología (CNB-CSIC), Campus de Cantoblanco, Madrid 28049, Spain.

## Abstract

By building on the SEVA (Standard European Vector Architecture) format we have refactored a number of regulatory nodes recruited from both Gram-negative and Gram-positive bacteria for rigorously comparing and parameterizing five expression devices that respond to diverse and unrelated chemical inducers, i.e. LacI^q^-P_trc_, XylS-P_m_, AlkS-P_alkB_, CprK-P_DB3_ and ChnR-P_chnB_. These were assembled as cargoes following the SEVA standard within exactly the same vector backbone and bearing the different functional segments arrayed in an invariable DNA scaffold. Their performance in an *Escherichia coli* strain of reference was then analyzed through the readout a fluorescence reporter gene that contained strictly identical translation signal elements in all cases and in the same DNA context. This study allowed us to describe and compare the cognate expression systems with unprecedented quantitative detail. The systems under scrutiny diverged considerably in their capacity, expression noise, inducibility and OFF/ON ratios. These features, along with the absence of physiological effects caused by the inducers and the lack of cross-regulation offer a panoply of choices to potential users and help interoperability of the specific constructs.

## INTRODUCTION

Prokaryotic transcription factors (TFs) responsive to exogenous physicochemical signals and their target promoters have been an invaluable source of biological parts for designing heterologous gene expression (Baneyx, 1999; Terpe, 2006; Yin *et al.,* 2007). Many of them rely on DNA sequences of the TF gene expressed through its own native promoter (typically a weak one) placed artificially close to the binding sites of the regulator adjacent to the site where transcription starts (Marschall, 2017). The literature contains a very large number of variants of such a basic theme, most often assembled in dedicated plasmid vectors that may be endowed also with additional features to optimize expression of the gene(s) of interest i.e. copy number, toxicity, mRNA stability, translation efficiency and other features (Zerbs *et al.,* 2009; Kaur *et al.,* 2018; Kent and Dixon, 2020). The vast majority of heterologous expression systems are based on *E. coli* as the host and TFs/promoters retrieved from regulatory networks of the same organism. Popular devices to this end include LacI/Plac promoter (Heyneker *et al.,* 1976; Lanzer and Bujard, 1988; Glascock and J. Weickert, 1998), TetR/PtetA (Skerra, 1994; Korpela *et al.,* 1998; Ramos *et al.,* 2005), AraC/ParaBAD (Guzman *et al.,* 1995; Haldimann *et al.,* 1998), RhaS/PrhaBAD (Egan and Schleif, 1993; Haldimann *et al.,* 1998), CI857/PL (Elvin *et al.,* 1990; Love *et al.,* 1996), which can be combined with the use of the extremely selective viral T7 RNAP/PT7 transcription machinery (Studier and Moffatt, 1986; Sørensen and Mortensen, 2005). Signals for triggering expression with these modules may vary from actual natural inducers to synthetic structural alternatives as well as mere physical cues e.g. temperature, cell density etc (Andrews *et al.,* 1996; Scholz *et al.,* 2003; Henßler *et al.,* 2005; Tang *et al.,* 2008; Choudhary and Schmidt-Dannert, 2010; Tang and Cirino, 2011; Kelly *et al.,* 2016; Taylor *et al.,* 2016; Ellefson *et al.,* 2018; Meyer *et al.,* 2019). In principle, these systems along with specialized strains have covered thus far most if—not all—necessities for engineering in *E. coli.* Yet, it is puzzling that despite the abundance and importance of such systems for contemporary biotechnology so little has been made to format them with a precise genetic architecture and rigorously describe them in a quantitative fashion. Fortunately, a growing number of expression devices have been inspected in recent years owing to the push for standardization brought about by synthetic biology (Canton *et al.,* 2008; Beal *et al.,* 2020). But for the most part, the best studied modules aimed at engineering complex expression circuits are largely limited to *E. coli.* In contrast, many other useful expression systems—specially those intended for non-model Gram-negative microorganisms—are poorly characterized and subject to all types of capricious designs and vector architectures. As a consequence, despite of the large number of potentially useful regulatory parts available for engineering heterologous gene expression (Calero *et al.,* 2016), a reliable metrics of their performance of most of them is lacking. Fortunately, recent developments are improving this state of affairs. Since 2013 there is an expanding collection of genetic tools (including expression systems) that follow the assembly rules of the so-called Standard European Vector Architecture (Silva-Rocha *et al.,* 2012), what enables reusability, portability and comparative assessment of promoter and genetic device performance in a variety of Gram-negative hosts (Tas *et al.,* 2021). This standard has been further expanded recently (Arce-Rodriguez *et al.,* 2020) to propose a specific architecture for heterologous expression modules consisting a fixed DNA sequence that acts as the scaffold for incorporation of the regulator gene and the target promoter, all organized in an otherwise invariable array of segments within a SEVA vector frame. The value of this format for expression cargoes was shown in the design and the superior performance of an improved module based on the AlkS/P_alkB_ regulatory node. This device included parts from the OCT plasmid of *Pseudomonas oleovorans* (van Beilen *et al.,* 2001; Rojo, 2009) and can be triggered by *n-*octane as well as by the gratuitous inducer dicyclopropyl ketone (DCPK).

In this work, we have expanded the standardization of expression modules to four additional platforms based on regulatory actors mined from Gram-negative bacteria in three cases, i.e. the *E. coli* (IPTG-inducible LacI^q^-P_trc_; Brosius *et al.,* 1985), *Pseudomonas putida* (XylS/P_m_ that responds to benzoate and *m-*toluate)(Gawin *et al.,* 2017) and *Acinetobacter* sp. (the cyclohexanone-responsive ChnR/P_chnB_ system; (Steigedal and Valla, 2008). In addition, one more expression system originated in the regulatory node CprK/P_DB3_ of Gram-positive organism *Desulfitobacterium hafniense*, which can be induced by chlorinated intermediates such as 3-chloro-4-hydrophenylacetic acid CHPA (Kemp *et al.,* 2013). In order to make them comply with the SEVA format, the biological parts employed in this study were edited to erase undesired restriction sites that could interfere with cloning and debugged to the maximum in order to improve their functioning, without affecting their functionality. As explained below, the key parameters that rule the performance of each system could then be rigorously determined and the data evaluated for fitting specific bioengineering goals. Although each of the pieces chosen for this study came from a different biological context, their formatting in a comparable genetic arrangement permitted a faithful appraisal of their duty in a given host. While *Pseudomonas* and other environmental bacteria are a virtually inexhaustible source of regulatory parts for engineering heterologous expression (Nikel and de Lorenzo, 2018), the work below with just 5 modules showcases the value of standardization of such devices and paves the way to a much ampler repertoire of choices available for designing complex bacterial phenotypes.

## RESULTS AND DISCUSSION

### Rationale for formatting SEVA-compatible heterologous expression cargoes

The 5 expression systems under examination in this work include LacI^q^-P_trc_; XylS/P_m_, ChnR/P_chnB_, AlkS/PalkB and CprK/P_DB3_. The components of each of them were edited and assembled to meet SEVA standards (Silva-Rocha *et al.,* 2012), so that all share the following common features [i] a constitutive promoter driving the expression of [ii] the gene coding for the regulatory protein, [iii] a spacer sequence (buffer) that insulates the expression from the adjacent DNA features and [iv] the cognate promoter that is activated upon addition of the inducer molecule. All the expression systems were flanked by SEVA-compatible PacI-AvrII sites, allowing easy swapping between SEVA plasmids. Downstream the regulator/promoter system there is a multiple cloning site (MCS) that facilitates the insertion of genes of interest, followed by a universal M13 (F24) priming site for sequencing/checking purposes. The functional cargo ends in a unique SpeI site so that, once inserted into any SEVA vector as a PacI-SpeI DNA segment, the expression cargoes described above are then encompassed by the strong terminators T0 and T1, which assure their transcriptional insulation (Fig. 1, upper panel).

**Figure 1.**
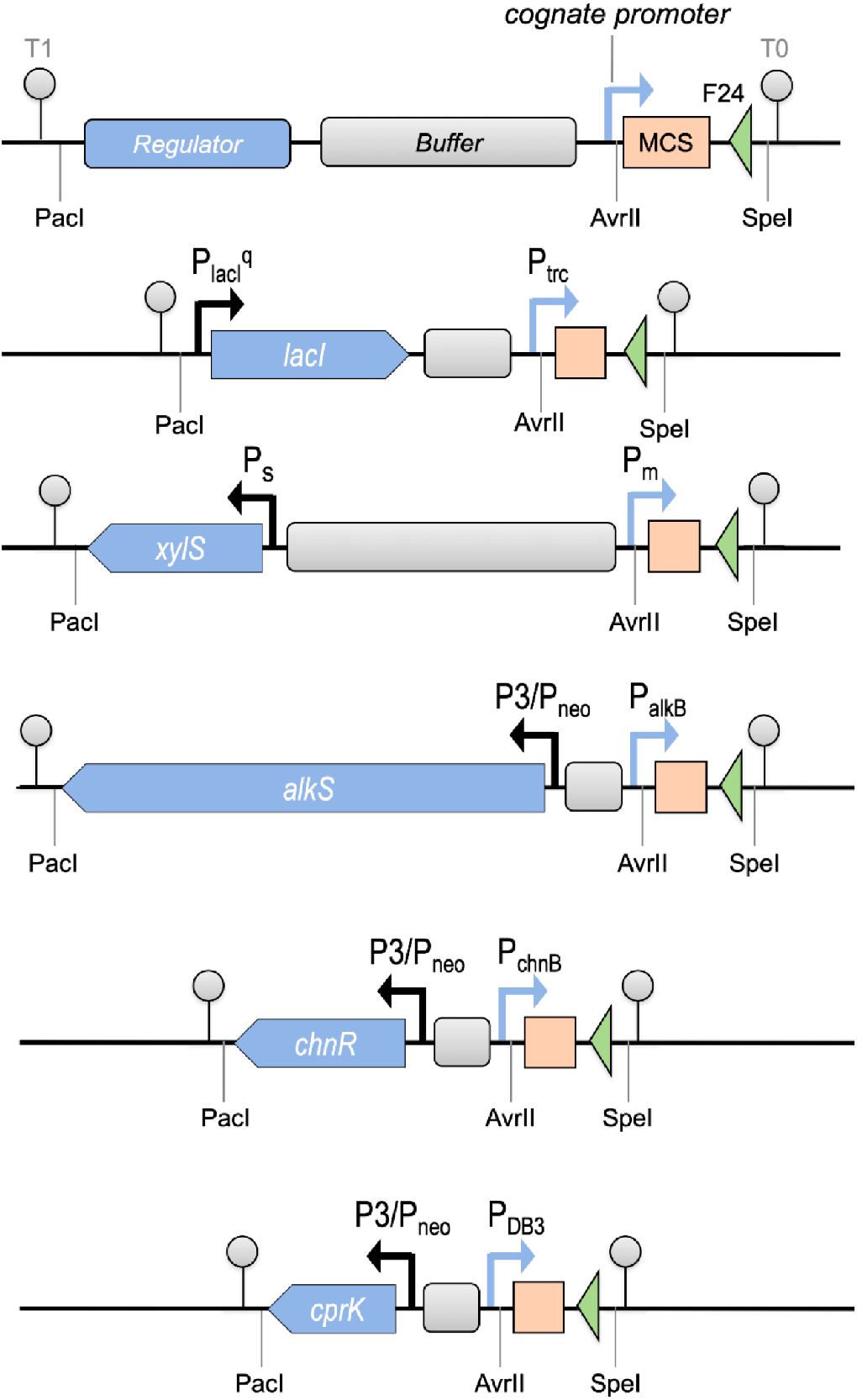
Main features of the standardized expression systems The upper panel shows a schematic representation of the main features encompassed within a pSEVA-compatible expression cargo, indicating the relevant elements involved in the regulatory device, i.e. transcriptional regulator, cognate promoter and a buffer sequence that separates both features. A handy MCS for cloning procedures locates downstream the regulatory device. In addition, there is a F24 oligonucleoitde priming site that can be used for checking/sequencing purposes. The unique restriction sites PacI, AvrII and SpeI allow modules exchange within SEVA platform backbones, in such a way that the whole module be flanked by appropriate transcriptional terminators T0 and T1. The structure of the five expression cargoes have been depicted to scale in the inserts below in the following order (from top to bottom): LacI^q^-P_trc_, XylS/P_m_, AlkS/P_alkB_, ChnR/P_chnB_, CprK/P_DB3_. Note that LacI^q^-P_trc_ and XylS/P_m_ have different spacers and specific promoters driving the expression of the corresponding TF, while the latter three expression systems, i.e. AlkS/P_alkB_, ChnR/P_chnB_ and CprK/P_DB3_ share a complete standardization of the region between the transcription factor and its cognate promoter, as explained in the text

The transcription factors (TF) locate divergently with respect to their cognate promoters in all cases with the exception of the IPTG-inducible system, were the gene encoding the regulatory protein LacI share the same orientation as its cognate P_trc_ promoter (see Fig. 1). As mentioned above, a DNA spacer separates the regulatory protein from its target promoter in all cases. This neutral DNA segment, which has no known regulatory elements, acts as a buffer sequence that alleviates the negative supercoiling originated from the transcription of the flanking promoters (Wu *et al.,* 1988). In three cases, i.e. AlkS/P_alkB_ (Arce-Rodriguez *et al.,* 2020), ChnR/P_chnB_ (Benedetti *et al.,* 2016) and CprK/P_DB3_ (Benedetti, 2014), the DNA buffer sequence has been standardized and thus it has identical length and sequence in all constructs, as described. In addition, in all these three expression devices a P3/P_neo_ promoter and a translation initiation region (TIR) drive the constitutive production of the regulatory proteins AlkS, ChnR and CprK, respectively. An additional edit was done in the *alkS* gene were a Crc recognition site was removed to mitigate the catabolic repression mediated by some carbon and nitrogen sources, exerted through a post-transcriptional signal recognized by the Crc/Hfq complex of *Pseudomonas* (Hernández-Arranz *et al.,* 2016); see (Arce-Rodriguez *et al.,* 2020) for more details on the standardization of AlkS/P_alkB_ device. In contrast, XylS/P_m_ and LacI^q^-P_trc_ pairs, whose sequences were obtained from well documented sources, were separated by spacers of different length and nature (Fig. 1; Silva-Rocha *et al.,* 2012). In any case, these buffer DNA sequences are intended for regulator/promoter insulation and not much impact of those small differences (if any) on the functionality of the device is expected.

Another feature that differs in the case of XylS/P_m_ and LacI^q^-P_trc_ expression systems is that the production of the respective regulatory protein is controlled by their own native constitutive promoter, in both cases. Concerning the IPTG-inducible system, the production of the TF LacI is driven by P_lacI_^q^, a mutant version of the P_lac_ natural promoter that leads to increased transcription and thus higher levels of repressor. This configuration was maintained to assure complete repression of the hybrid *trp/lac* cognate promoter (Calos, 1978; Amann *et al.,* 1988). With respect to XylS-P_m_ regulatory device, previous expression studies reported that in high copy-number plasmids the basal level of the Pm promoter increased more than in proportion to the copy number (Mermod *et al.,* 1987; Ramos *et al.,* 1990; Michan *et al.,* 1992; Kessler *et al.,* 1994), observations that seem to be related to the ability of the regulator XylS to activate P_m_ in the absence of inducer, provided it is present at high intracellular concentrations (Inouye *et al.,* 1987b; Ramos *et al.,* 1987; Spooner *et al.,* 1987). For this reason, we decided to keep the relatively weak native P_s_ promoter to maintain XylS levels low, in order to minimize leaky activity from the P_m_ promoter.

### Single-cell analysis of the expression dynamics by flow cytometry experiments

To prevent the occurrence of effects unrelated to the features of the regulation/promoter systems themselves, all replicons used in this study shared an identical backbone i.e. the plasmid pSEVA231, harboring a medium copy-number pBBR1 origin of replication and a kanamycin resistance gene (Silva-Rocha *et al.,* 2012). Following SEVA standard, the expression modules reported in the above section were in all cases inserted in exactly the same location, i.e. between the PacI-AvrII sites. The cloning procedures originated plasmids pSEVA234 (containing LacI^q^/P_trc_ features), pSEVA238 (harboring XylS/P_m_ cassette), pSEVA239 (carrying the regulator/promoter regions of AlkS/P_alkB_), pSEVA2311 (bearing ChnR/P_chnB_ regulatory pair) and pSEVA2312 (for CprK/P_DB3_ expression system). A reporter gene coding for the monomeric and superfolder GFP (msfGFP), bearing a TIR motif (Miller and Lindow, 1997), was then cloned downstream all the regulator/promoter systems under study (see experimental procedures) at the same position of the MCS, in order to monitor, quantify and compare their activity. The resulting constructs named pS234M, pS238M, pS239M, pS2311M and pS2312M, harboring LacI^q^-P_trc_, XylS/P_m_, AlkS/P_alkB_, ChnR/P_chnB_ and CprK/P_DB3_ expression systems, respectively, which were used in the following experiments. In all instances *E. coli* CC118 was used as host strain that was grown under equivalent conditions, as a means to minimize uncontrolled genetic and cellular context influences. The first aim of this work was to compare general performance of these expression systems in terms of growth, inducibility and production yield. The analysis of the GFP production driven by each of the five expression cassettes described above was done by means of flow cytometry. In contrast to the experiments carried out at the level of cell population, which obscure significant differences in expression levels of individual cells, flow cytometry brings to light such cell-to-cell heterogeneity and therefore provides a more detailed information. Fig. 2 shows the fluorescence patterns of cells harboring the expression system under study, along time after addition of saturating amounts (1 mM) of the corresponding inducer, as indicated. A *E. coli* CC118 host carrying the plasmid pS237M, a promotor-less msfGFP gene inserted in the same backbone as the constructs under analysis, was used as non-expressing control.

**Figure 2.**
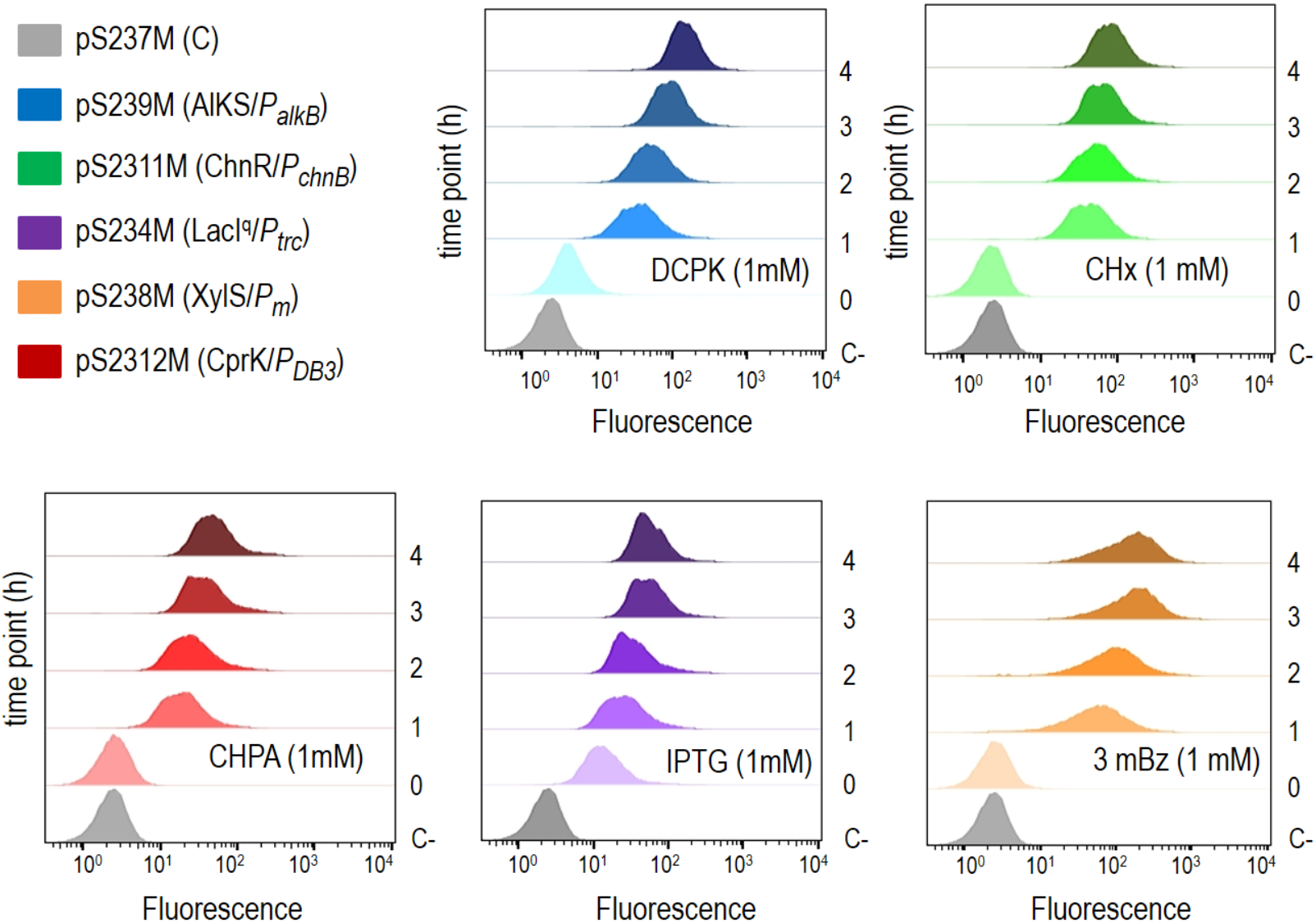
Analysis of the fluorescence distribution using flow cytometry *E.coli* CC118 standard laboratory strain harboring the corresponding expression plasmids, as indicated, were grown under standard conditions up to the mid-exponential phase (OD≈0.4) when the cultures were induced with the corresponding effectors (t=0). Samples were collected at one hour frequency after induction (t=1, 2, 3 and 4 hours), fixed and the GFP signal collectively analyzed with a flow cytometer. Each histogram is one representative sample.

A first inspection of the frequency distribution histograms for green fluorescent cells revealed significant differences among the regulator/promoter systems used in this work. In the absence of any inducer, the populations remained mostly non-fluorescence in any case but in the LacI^q^-P_trc_ expression system, were a significant percentage of cells showed a detectable GFP signal, indicating a relatively high basal level. As expected, all the expression systems responded to the addition of the corresponding inducer, as reflected by the fluorescence signals that moved towards higher intensity values over time. In all cases, fluorescence values fell in a single peak but the histograms showed diverse shapes and fluorescence accumulated with different dynamics, depending on the time-point and the expression system under analysis. Four hours post induction was enough to reach a plateau of accumulated GFP fluorescence in all the expression systems under the conditions of the analysis.

### OFF/ON expression levels among regulator/promoter systems differed to a great extend

A comprehensive analysis of the results obtained by means of the flow cytometry experiments described in the previous section allowed as to draw interesting conclusions about the behavior of the expression systems under consideration, both in the OFF and in the ON states. First, we had a look and compare uninduced production levels. The tightness of gene regulation is an important quality feature for expression systems. Even low basal activity can have adverse or toxic effects on the host metabolism, what makes very challenging or even impossible the heterologous expression of certain specific proteins. Moreover, circuit-associated burden is an important consideration that must be accounted for engineering genetic circuits of increasing complexity. In addition, to have the lowest possible background—ideally no expression in the OFF state— is essential to build circuits in a predictable manner, allowing timely induction as required. Furthremore tight regulation of basal expression enables handling of toxic genes.

A detailed analysis of the fluorescence values obtained at mid-exponential growth and before adding the corresponding inducer (time point t=0) revealed important differences among the median values of fluorescence basal levels (Fig. 3). LacI^q^-P_trc_ was, by far, the leakiest system, producing around 55 times more GFP protein than the ChnR/P_chnB_ regulator/promoter system, which behaved as the tightest device under non-inducing conditions (fluorescence median of 989.5 and 17.5, respectively), followed by XylS/P_m_ and CprK/P_DB3_ both displaying similar basal levels (fluorescence median of 53.5 and 43, respectively), which roughly doubled that of the ChnR/P_chnB_ expression system. The regulatory stringency was intermediate in AlkS/P_alkB_ (fluorescence median of 221.5 a.u.) that exhibited five times less background than the LacI-dependent system but was between 4 and 12-fold leakier than the rest of expression systems.

**Figure 3.**
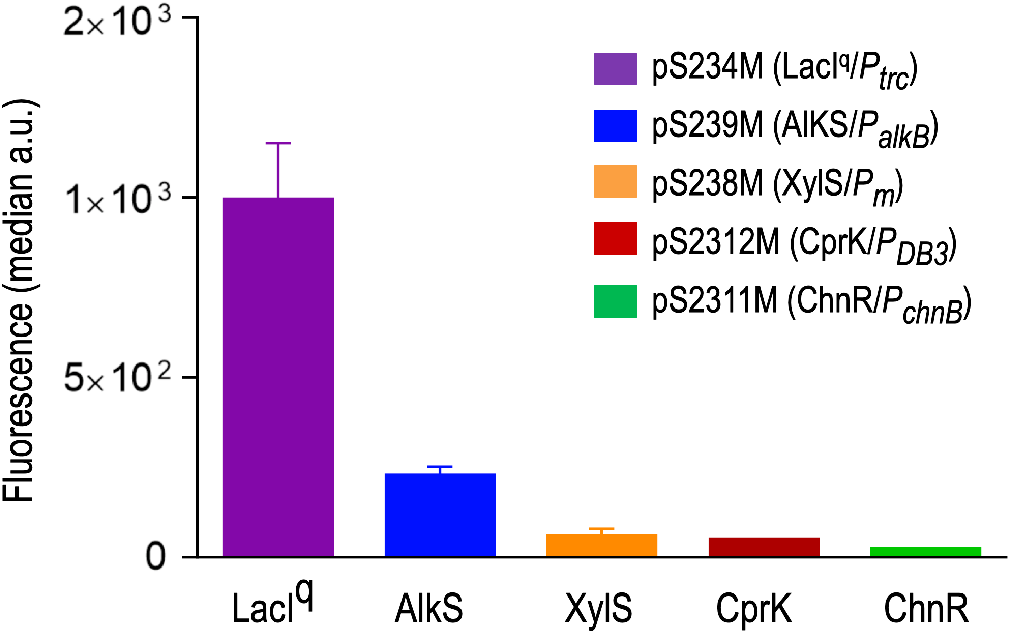
Tightness of regulatory/promoter systems Basal expression of *E. coli* CC118 cells harboring the corresponding expression plasmids (t=0), were obtained from the median values of flow cytometer experiments and plotted for a comparison, as indicated. Values correspond to three independent experiments with two technical replicates each. Basal expression levels were ordered from highest to lowest below the diagram. By far, the leakiest system was LacI^q^-P_trc_.

The P_trc_ core promoter is a chimaera of the *E. coli trp* operon and lac UV5 promoters, including a single proximal lacO1 from the lac operon and the optimally spaced *E.coli* consensus −35 and −10 motifs. In the absence of lactose (or IPTG, a non-metabolizable structural mimic of allolactose) the lac repressor protein LacI binds the operator site (lacO) preventing transcription from the P_trc_ promoter. The observation of an inefficient repression or P_trc_ in the conditions tested could be due to a suboptimal ratio between LacO sites with respect to the amount of LacI repressor molecules. In negatively regulated expression systems such as LacI and TerR-related layouts, it is well stablished that the higher the number of operators contained, the lower the basal expression is (Oehler *et al.,* 1990; Oehler *et al.,* 1994; Bertram and Hillen, 2008). Therefore, it is likely that both an increase in the number of LacI molecules and/or the addition of a second lacO site might reduce the basal expression. However, increasing the strength of the repression usually results in reduced transcriptional activity under inducing conditions too. An excessive amount of operators could lead to almost complete stop of production even in the activated state and in consequence the number of *lacO* operator sites must be well balance (Schuller *et al.,* 2020). Another reason that could account for the remarkably high basal level of gene expression in this case could be the potential readthrough transcription originated in the *lacI* gene that is located upstream and in the same orientation as its P_trc_ target promoter. It is likely that placing the TF in a divergent orientation with respect to its target promoter would prevent pervasive transcription and therefore decrease the basal expression.

Outstanding differences were also observed among the regulatory/promoter devices with respect to their transcriptional capacity at the activated ON state (see Fig. 4A). In all cases, GFP fluorescence accumulated steadily along time, upon addition of the corresponding inducer. However, the maximum protein yield -after four hours of induction- largely diverged depending on the expression system. The XylS/P_m_ cassette reached the highest GFP production followed at a short distance by the AlkS/P_alkB_ system. In contrast, the lowest protein producers were CprK/P_DB3_ and LacI^q^-P_trc_ expression devices, which displayed up to four times less GFP fluorescence. Finally, ChnR/P_chnB_ showed around half the amount of GFP production compared to the maximum rates exhibited by the best performers XylS/P_m_ and AlkS/P_alkB_ expression systems. Thus, in terms of absolute reporter output the yield ranking was established as follows: XylS/P_m_ > AlkS/P_alkB_ > ChnR/P_chnB_ > CprK/P_DB3_ = LacI^q^-P_trc_.

**Figure 4.**
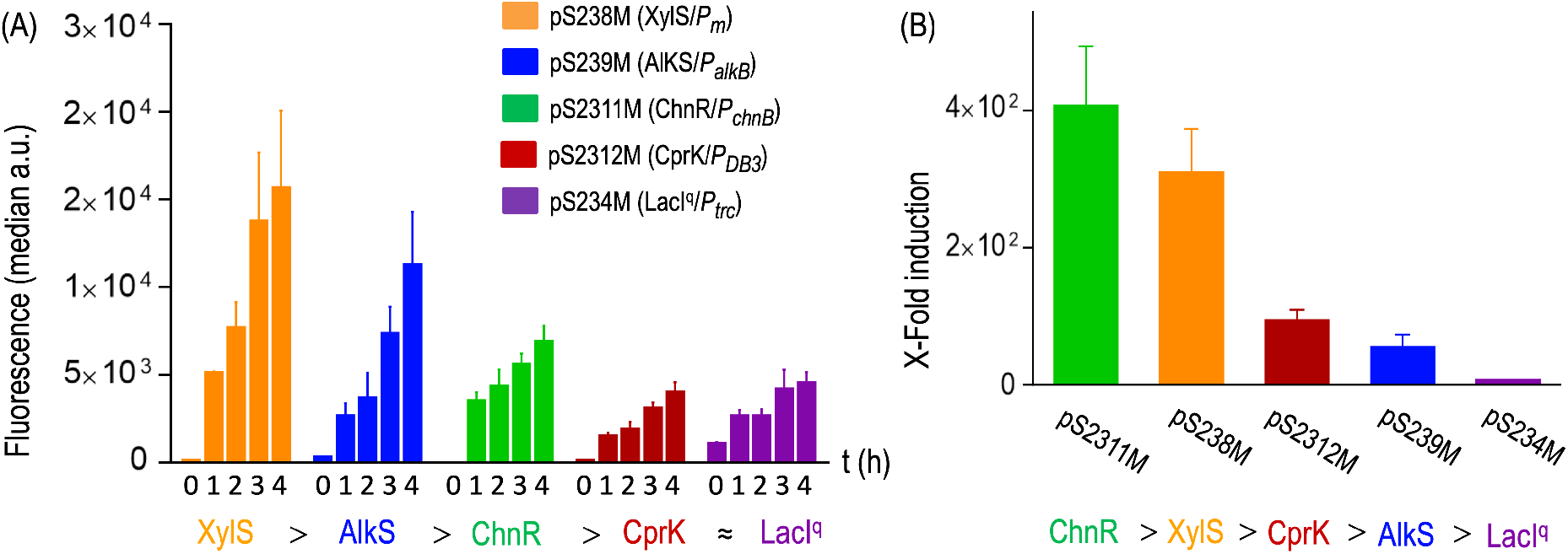
Comparative activated expression along time and dynamic ranges (A) Median values of the activated expression of cultures of *E. coli* CC118 cells carrying the corresponding expression plasmids, obtained from three independent flow cytometer experiments, were plotted as for comparison of their expression levels along time (t=0, before induction; t=1, 2,3 and 4 h after adding the inducer, respectively). The highest expression values, reached at 4 h after induction, served to order the production capacity of each regulatory device, which are indicated below the graph in descending order. (B) X-fold induction of the different expression systems obtained from flow cytometry experiments, as described. Rank from the highest to the lowest dynamic range among the regulatory devices is outlined below the plot.

All in all, the dynamic range of the system, i.e. the production ratio between activated and non-induced expression levels (also defined as induction ratio or induction window), provides a most interesting parameter of a given system behavior. In addition, using the dynamic range criterion facilitates the comparison of data obtained from various sources. As shown in Fig. 4B, ChnR regulated device displayed the highest induction ratio with 388-fold protein production after induction (green bar). Another expression system that exhibited a good dynamic range was Xyl/P_m_ (Fig. 4B, orange bar), which increased GFP production by 291-fold when induced. CprK-dependent and AlkS-regulated expression systems showed relatively good induction rates too (91-fold and 50-fold changes, respectively) while the LacI^q^-P_trc_ was by far the least inducible system as GFP fluorescence increased barely 5-fold upon induction. Taking into account the dynamic range as a good descriptor of the expression capabilities, the regulatory devices analyzed in this work rank as follows (from best to worse): ChnR/P_chnB_ >XylS/P_m_ > CprK/P_DB3_ > AlkS/P_alkB_ > LacI^q^-P_trc_

### Gene expression noise: phenotypic diversification

Flow cytometry profiles embody valuable information about cell-to-cell homogeneity in a given population, in contrast to bulk fluorescence analysis, in which the signal is measured as the population mean fluorescence, masking important differences in the level of expression among cells that are known to occur. The width of the flow cytometry peaks is directly proportional to the spread of the individuals within the population. Therefore, a sharp peak indicates low dispersion or higher cell-to-cell homogeneity while a wider peak indicates higher variation within the expression level among individual cells. A good indicator of the level of heterogeneity is the coefficient of variation (CV), a parameter that relates the standard deviation with the mean value and is usually expressed as a percentage. In practical terms, the lower the CV the less the noise there is. As CV is unitless and dimensionless, this proxy can be used to compare experimental results over time as well as those obtained in different platforms.

Fig. 6 shows the CVs (expressed as percentage), of *E. coli* CC118 strains harboring the corresponding expression systems before and at different time points after induction, as indicated. At time point 0, right before adding each inducer, most of the expression systems had a low spread, i.e. a small CV that was around 45 to 50%, with the exception of the LacI^q^-P_trc_ regulatory system that displayed a mean CV of 63%, representing the highest heterogeneous fluorescence signal among cells at the uninduced estate. This intrinsic noisy behavior is most likely due to the stochastic expression due to fluctuations arising from the suboptimal repression of the P_trc_ promoter exerted by the LacI regulator, what translates into a broader expression profile. This hypothesis is supported by the high basal expression displayed by this expression system, as indicated previously.

**Figure 5.**
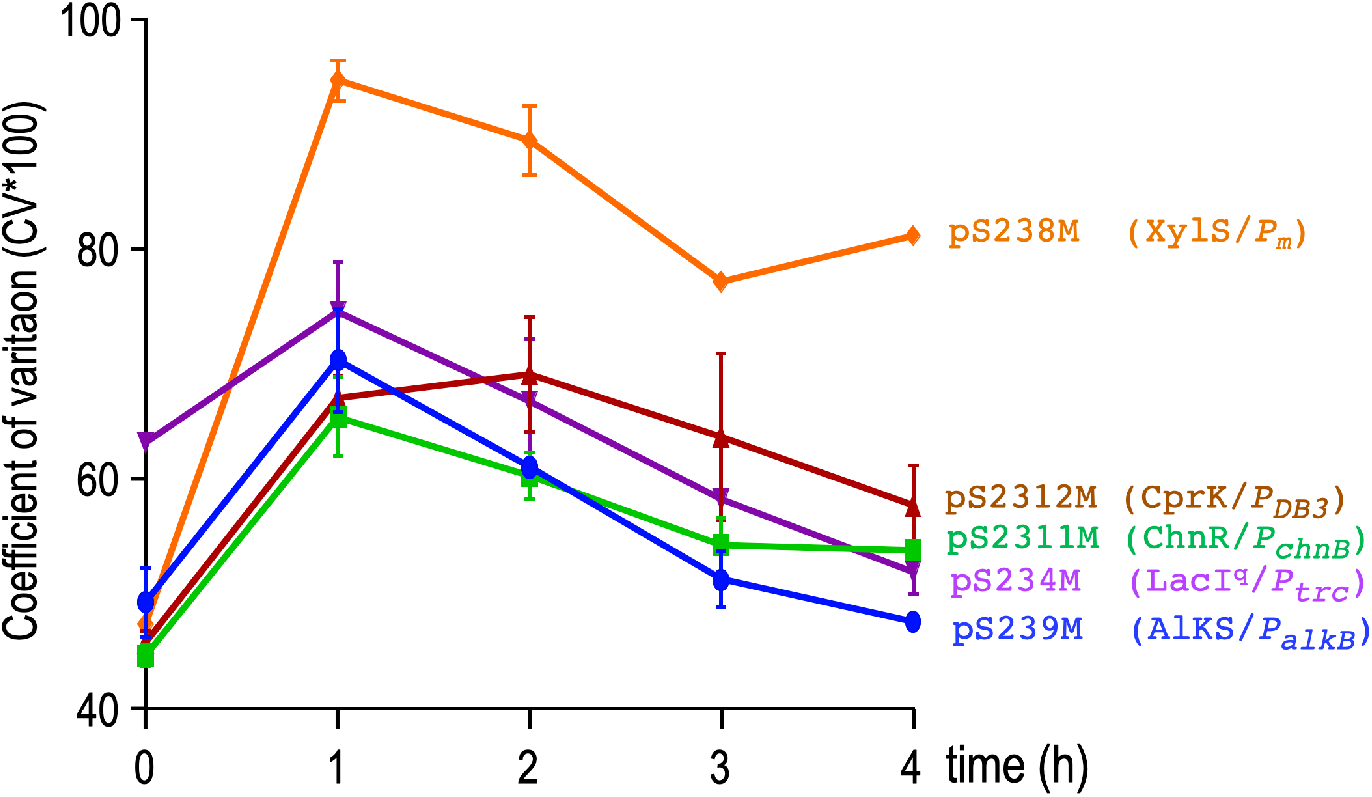
Population heterogeneity Representation of the cell-to-cell heterogeneity of *E. coli* CC118 populations harboring the corresponding expression systems along time, expressed as percentage of the coefficient of variation (CV*100) at each time point of the flow cytometry experiments (t=0 before induction, t=1, 2, 3 and 4 h after addition of the inducers). Below the diagram the list of expression systems have been ordered with respect to their noise in descending order (the noisest system is XylS/P_m_ while the ones with the lowest cell-to-cell heterogeneity are CprK/P_DB3_ and AlkS/P_alkB_).

**Figure 6.**
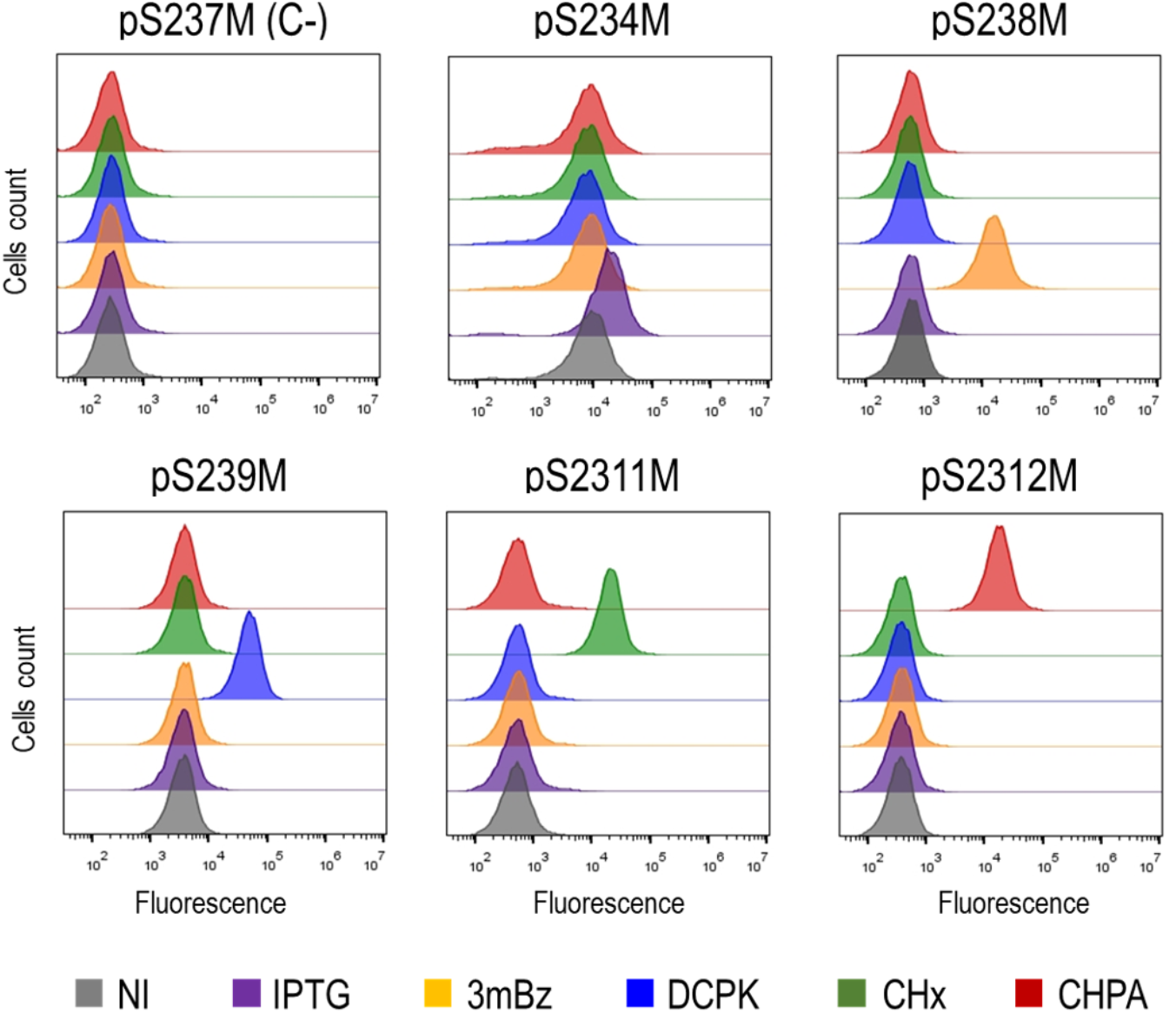
Cross-talk among the expression systems *E. coli* CC118 cells transformed with the corresponding expression plasmids, as indicated above each graph, were either non treated (NI) or induced with their own effector as well as with the rest of the heterologous inducers involved in the expression systems under analysis, i.e. IPTG (orange color), 3mBz (blue color), DCPK (red color), CHx (green color) and CHPA (purple color). The effect of the corresponding induction on the GFP fluorescence was then analyzed by flow cytometry experiments after two hours of treatment, as indicated. A host strain harboring the promotorless msfGFP gene (pSEVA237M) was also tested as negative control. Note that the expression systems responded only to their cognate induces and not to the effectors activating other regulators.

Upon activation by their cognate inducers the cell-to-cell fluorescence variations increased in all the expression systems. At short time of analysis, i.e at 1 hour post induction, CV values exceeded 60% in all samples, but decreased steadily over time. In all cases except for the XylS/P_m_ expression system, after four hours of induction population cell-to-cell homogeneity was similar to that observed at the beginning of the experiment. The XylS/ P_m_ regulator/promoter system showed the most uneven signal distribution at the ON estate throughout the whole period, especially at early times post-induction, as reflected by a CV value that reached almost 100% dispersion, i.e. the standard deviation was similar to the mean vale one hour after the addition of 3mBz. Although the spread of XylS-dependent expressing cells decreased over time, it remained higher than any of the rest of expression systems under study was, and came to around 80% at late times post-induction. In general, fluctuations in GFP expression observed in this piece of work are consistent with values previously reported for similar expression systems (Balzer *et al.,* 2013). Gene expression noise arises from two main components, i.e. intrinsic noise (transcriptional and translational bursts) and extrinsic noise (general fluctuations on cell components) and thus cell-to-cell variation in GFP expression is affected from both types of variation (Elowitz *et al.,* 2002; Swain *et al.,* 2002; Paulsson, 2004; Kærn *et al.,* 2005; Lestas *et al.,* 2008; Hilfinger and Paulsson, 2011). However, a major contribution of stochasticity can be traced to the dynamics of the transcription factor (TF)-promoter interplay (Chalancon *et al.,* 2012; Munsky *et al.,* 2012). A significant component of the gene expression noise can be attributed to the average protein abundance, leading to a phenomenon called noise propagation, i.e. the expression noise within a specific gene can result from fluctuations in levels of its cognate TF (or of the signaling molecules, influencing the TF activity; Pedraza and van Oudenaarden, 2005; Bar-Even *et al.,* 2006; Sanchez *et al.,* 2013). Thus, the noisiest behavior observed in XylS/P_m_ regulatory system could be explained, at least in part, by the relative low protein abundance of XylS. In its natural context, i.e. the TOL plasmid pWWO, XylS is overproduced via the XylR master regulator that also activates intracellular 3mBz production, what leads to high levels of active XylS regulator (Inouye *et al.,* 1987a,b; Marqués and Ramos, 1993). Here, XylS production is driven by the basal expression from the Ps promoter at limited amounts. It has been reported that low levels and/or a low availability of XylS translates into increased P_m_ promoter noise. Modelling of XylS/P_m_ interplay indicated that the more XylS active form the less noisy P_m_ behaves. This prediction was supported by the observation that a P_m_ promoter located in trans with respect to its cognate regulator XylS, at a distance that decreased the number of active XylS TFs at the local molecular proximity of the promoter, showed a more noisy performance than an equivalent XylS/P_m_ assembly placed in *cis* (Goñi-Moreno *et al.,* 2017).

### Cross-regulation among expression systems

The size and complexity of gene networks is often limited by the amount of regulatory parts that can be used without interference. Orthogonality among expression systems is needed to eliminate undesirable crosstalk and to allow a separate dynamic control of expression modules. Therefore, we next analyzed potential crosstalk effects from effectors others than the specific one of any of the regulatory devices under analysis. To do so, an *E. coli* CC118 host strains harboring the corresponding expression system were separately supplemented with each of the different inducers under consideration (or non-treated), as indicated. A control strain, carrying the promotorless GFP plasmid pS237M was also included in the assay. After three hours of treatment, the effect on GFP fluorescence was monitored by means of flow cytometry experiments. As shown in Fig. 6 each expression system responded to their inducer, displaying a shift of the population toward higher fluorescence values, as expected. However, none of the inducers affected neither the control nor the strains carrying plasmids with heterologous expression systems, which showed similar pattern of fluorescence as those of the corresponding untreated sample. Potential toxic effects of inducers were also tested by means of growth curve of cells endowed with each of the expression systems that were separately treated with the whole array of inducers. Growth patterns were identical in all cases to that of the corresponding untreated sample, suggesting that none of the chemicals had a significant impact in cell physiology (not shown). These experiments thus indicated that the expression systems reported in this work are not influenced by inducers other than their cognate effector.

### Inducers-mediated release of Reactive Oxygen Species (ROS)

To further characterize the expression systems under analysis we next sought to see if any of the inducers involved in activating the corresponding regulatory proteins were able to cause oxidative stress in the host strain used in this study. Various stressors might exert their toxic effects by amplifying the natural rate of reactive oxygen species (ROS), which can damage vital molecules inside the cell like DNA, membranes and proteins. It also alters the redox status inside the cell, leading to pleiotropic effects that could even determine cell death (reviewed in Cabiscol *et al.,* 2000; Zhao and Drlica, 2014). Potential toxic effects and release of oxidative stress related compounds, as consequence of the addition of the inducers, were investigated by means of growth curves and direct quantification of ROS by flow cytometry (Fig. 7). As shown in Fig. 7A, the addition of inducers at the concentration employed in this work (1mM) did not inhibit growth of *E. coli*. Neither the growth rate nor the final optical density were affected by the addition of any inducer compared to the growth pattern of a non-treated control strain. In contrast, hydrogen peroxide (H_2_O_2_) and diamide worked as bacteriostatic agents in cells exposed to both chemicals, leading to growth arrest at approximately one hour after addition. Diamide, which acts as a drainer of metabolic NADPH, causes thiol crosslinking while H_2_O_2_ is the source of hydroxyl radicals that are the origin of relevant oxidative damage. Since both molecules are known to be ROS generating stressors, they were used as positive control in the experiment. Flow cytometry using 5-(6)-carboxy-2’,7’-dichlorodihydrofluorescein diacetate (carboxy-H2DCFDA) as the ROS-sensitive probe (Fig. 7B) revealed that none of the inducers produced a detectable increase in fluorescence, indicating that their oxidative effects were negligible. On the other hand, addition of both H_2_O_2_ (1.5 and 5 mM concentration) as well as diamide (250 ◻M) produced a substantial effect leading to a robust fluorescence induction in the cells (>98% at the lowest concentration) under similar conditions.

**Figure 7.**
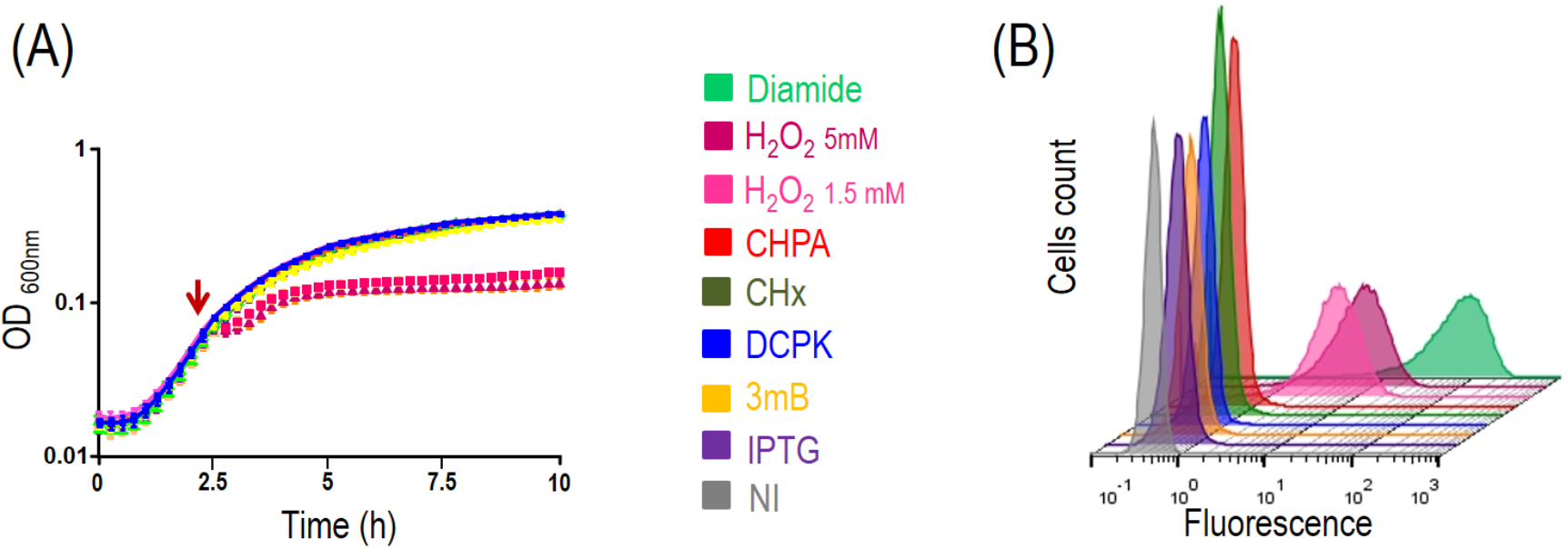
Inducer-mediated oxidative damage (ROS) (A) Growth curves of *E. coli* CC118 cells harboring plasmid pS237M treated with different inducers (at 1mM concentration) depicted with the same color code as in Fig. 6, or with ROS-producing agents, i.e. H_2_O_2_ at 1.5 and 5 mM concentration (soft and dark pink color, respectively) and diamide at 250 μM concentration (dark green color), as indicated. Addition of the different chemicals was done at mid-exponential growth (t=2.5 h, indicated with a red arrow). Non-treated cells (NI) were included as a control. (B) Flow cytometry experiments with samples of *E. coli* CC118 cells carrying plasmid pS237M treated as described above (see also experimental procedures). Two hours after the treatment, cell aliquots were analyzed using the oxidative probe carboxy-H2DCFDA. This nonfluorescent molecule is readily converted to a green-fluorescent form when the acetate groups are removed by intracellular esterases and oxidation (by the activity of ROS) occurs within the cell. The color code of each histogram, which are representative samples, corresponds to that of the growth curves in (A).

Altogether, the experiments shown in this section indicated that treatment of cells with any of the inducer connected with the expression systems considered here neither affected growth nor resulted in ROS generation in *E. coli* and therefore they could be considered as safe agents in this context.

## Conclusion

The application of a synthetic biology engineering mind-set has led to a significant increase in diverse and innovative gene regulation tools. One has to keep in mind which characteristics are best for a given application. For example, appropriate genetic control enables tight regulation. Other desirable features of expression systems are a wide dynamic range, tunable control of expression rate, high expression levels or robust performance. However, in spite of the impressive amount of useful expression systems available to date there is still a lack of reliable quantitative information about most of them. A further movement towards standardized measurements would allow direct comparison of performance, thereby enabling a robust selection of the most appropriate tools and expression modules for the construction of complex circuits. This need was the main motivation for this piece of work in which five SEVA-compatible expression cargoes have been first streamlined and next characterized. We determined and compared their relevant parameters, i.e basal and activated activity, population homogeneity and orthogonality. In addition, we ruled out the possibility of potential inducer-mediated toxicity. By capitalizing upon the wealth of regulatory components described here, which can be used stand alone or in combination, gene expression can be more precisely adjusted for a desired output. We thus expect the work presented here showcases the value of standardization and helps to widen the toolbox for the rational design of genetic networks.

## EXPERIMENTAL PROCEDURES

### Strains, plasmids, media and growth conditions

The relevant properties of the strains and plasmids used in this work are listed in Supplementary Table S1. *E. coli* DH10B, DH5α and CC118 strains were used for cloning procedures. *E. coli* CC118 strain harboring the corresponding plasmid under analysis, was used for assessing GFP activity driven by the different expression systems, as indicated. Bacteria were grown routinely at 37 °C in LB (10 g l^−1^ of tryptone, 5 g l^−1^ of yeast extract and 5 g l^−1^ of NaCl). When required, Km (75 ◻g/ml) was added to the culture media. The necessary inducers isopropyl-β-D-thiogalactopyranoside (IPTG), 3-methil-benozate (3mBz), cyclohexanone (CHx), dicyclopropyl ketone (DCPK) or 3-chloro-4-hydroxyphenylacetic acid (CHPA), were added where indicated to a final concentration of 1 mM. DNA synthesis was performed by GeneArt® (Thermo Fisher Scientific). Primers used in this work were provided by Sigma-Aldrich Co (St. Louis, MO, USA). Restriction enzymes and Phusion DNA polymerase used for PCR reactions were purchased from New England BioLabs inc. (Beverly, MA, USA). DNA AmpliTools Green Master Mix (Biotools, B & M Labs, SA Madrid-Spain) was routinely used for colony PCR verifications. The fidelity of DNA fragments was later confirmed by restriction mapping and DNA sequencing of relevant features (Secugen Sequencing and Molecular Diagnostics, Madrid, Spain).

### DNA constructs

General methods for DNA manipulation were performed as described (Sambrook, 1989). Oligonucleotides used in polymerase chain reaction experiments (PCR) are listed in Supplementary Table S2. The ITPG inducible expression system was obtained from the plasmid pTrc99a. The expression cassette containing the lacIq gene and the P_trc_ promoter was amplified by PCR using primers TRC-F and TRC-R These primers introduced the PacI and the AvrII restriction sequences in the flaking region of the fragment. A 1.4 kb PCR fragment was gel purified and cloned in the pSEVA221 vector previously digested with PacI/AvrII, leading to plasmid pSEVA224. The benzoate inducible XylS-Pm expression system was constructed based on the cassette found in the vector pJB653 and relatives (Blatny *et al.,* 1997). In that system, the XylS activator is placed in divergent position relative to its target Pm. As the XylS-Pm fragment presented many restriction sites used in SEVA multi-cloning site, the cassette sequence was edited in silico and the resultant variant was synthetized de novo by GeneArt. The 2.0 kb DNA fragment with the edited XylS-P_m_ sequence was cloned as a PacI/AvrII fragment into the pSEVA226 reporter vector, generating the plasmid pSEVA226-XylS-Pm, which was used to produce pSEVA238, as described (Calles *et al.,* 2019). The AlkS-P_alkB_ expression cassette -based in *P. putida* GPo1, was fully synthetized by GeneArt in order to fulfill SEVA standards and was finally cloned in pSEVA421 to produce pSEVA429, as described (Arce-Rodriguez *et al.,* 2020). Then, the PacI-AvrII fragment, containing the complete regulatory device, was recloned in pSEVA231 digested with the same enzymes, leading to plasmid pSEVA239. The design and construction of a standardized vector carrying the ChnR/P_chnB_ expression system is described elsewhere (Benedetti *et al.,* 2016). The CprK/P_DB3_ expression system was assembled as follows (Benedetti, 2014). The cprK1 gene coding sequence was amplified form plasmid pMLK1 (Kemp *et al.,* 2013), by a PCR reaction containing primers 5-CPRK1 and 3-CPRK1, harboring AvrII and PacI sites, respectively. The resulting DNA fragment was then digested with both enzymes and cloned into pSEVA231 sites, producing the intermediate auxiliary plasmid pSEVA231-CPRK1. A separate DNA fragment contained the cognate promoter P_DB3_, which was followed with a buffer sequence that separated from the divergent promoter P3/P_neo_, which was introduced along with the translation initiation region (TIR) sequence (5’-GATTAACTTTATAAGGAGGAAAAA-3’; Miller and Lindow, 1997), to drive the constitutive expression of the regulatory protein CprK. This fragment containing the DHPA inducible promoter plus the constitutive P3/P_neo_ promoter was cloned into NheI/AvrII sites of plasmid pSEVA231-CPRK1, producing plasmid pSEVA2312 that harbours the complete CprK/P_DB3_ expression system. The corresponding reporter plasmid pS2312M was obtained as indicated below. To obtain a plasmid for transcriptional fusions based in a promotorless msfGFP reporter we used plasmid pGA-LacI, containing an edited PstI-less version of the msfGFP gene, as template of a PCR reaction using oligonucleotides msfGFP HindII-F, which contained a TIR motif and a HindIII site, and msfGFP-SpeI, bearing a SpeI restriction site. The amplified fragment was then digested with HindIII and SpeI enzymes, gel purified and ligated into the corresponding sites of pSEVA 231, producing plasmid pSEVA237M (Benedetti *et al.,* 2016; Calles *et al.,* 2019). All plasmids used to monitor expression activity form the different systems, i.e. pS234M, pS238M, pS239M, pS2311M and pS2312M, were constructed by restriction of pS237M with enzymes HindIII-SpeI and ligation into the corresponding expression system previously cleaved within the same sites.

### Flow cytometry monitoring of msfGFP expression

For monitoring and quantifying nsfGFP expression in flow cytometry experiments, specific strains containing the relevant constructs, as described in each case, were inoculated into filtered LB medium and grown to stationary phase. Cells that had been grown overnight were then diluted to an OD_600_ of 0.05 in fresh filtered LB media and incubated at the appropriate temperatures to an OD_600_ of 0.4, when the corresponding inducer was added, as described. This time point was considered as t=0. Following exposure to the inducer, 1 ml samples were harvested at various time points, as indicated in the text, and spun down in a tabletop centrifuge at 13,000 rpm during 1 min; then, the cells were washed in 500 μl of filtered PBS 1X, centrifuged again as indicated above and resuspended in 300 μL of 0.4% (W/V) paraformaldehyde and incubated at room temperature for 10 min. After cell fixation, cells were washed twice in 500 μl of filtered PBS 1X and finally resuspended in 600 μL of filtered PBS 1X and stored on ice until analysis. The final OD_600_ was adjusted to a value < 0.4 in all samples before flow cytometry analysis.

Single-cell fluorescence of the samples was then analyzed by flow cytometry using a GALLIOS cytometer (Perkin Elmer) or with a MACSQuant™ VYB cytometer (Miltenyi Biotec, Bergisch Gladbach, Germany). GFP was excited at 488 nm, and the fluorescence signal was recovered with a 525/40 nm band pass filter. At least 25,000 events were analyzed for every aliquot. The GFP signal was quantified under the experimental conditions tested by firstly gating the cells in a side scatter against forward scatter plot, and then the GFP-associated fluorescence was recorded in the FL1 channel (515-545 nm). Data processing was performed using the FlowJoTM software as described elsewhere (www.flowjo.com). All experiments were carried out at least three times with two technical replicates.

### Flow cytometry detection of ROS (peroxides and hydroperoxides) by dichlorofluorescein derivatives

To detect ROS damage by means of flow cytometry analysis, a culture of *E. coli* CC118 cells transformed with pS237M was grown overnight as indicated. The following, samples were diluted in 25 mL of filtered LB to a OD_600_ of 0.05. At mid-exponential phase of growth (OD_600_ 0.4), the culture was divided into 7 mL aliquots that were treated either with the stressors H_2_O_2_ or diamide (as positive controls), as indicated or with 1mM of any of the inducers under inspections. One more aliquot was kept untreated as negative control. After two additional hours, 1 mL of the cultures was spun it down at 13,000×g for 2 min at room temperature, washed once with 1mL of filtered PBS 1X by centrifugation and resuspension and diluted in filtered PBS 1X such that OD_600_ was under 0.4 in a final volume of 1mL. Then, a freshly prepared solution of 5-(6)-carboxy-2’,7’-dichlorodihydrofluorescein diacetate (carboxy-H2DCFDA; Sigma-Aldrich Co., St. Louis, MO, USA) in DMSO, was added to the samples at a final concentration of 20 μM. Cells were incubated for 30 min at 37°C protected from light. Finally, the fluorescence of each sample was measured in the green channel using the excitation/emission parameters as for GFP.

### Growth curves

*E. coli* CC118 cells transformed with plasmid pS237M were grown overnight in LB medium at 37°C with routinely conditions described above and then diluted 1:20 in multiwell microtiter plates containing 190 μl LB medium, supplemented with kanamycin. Cells were the grown with rotatory agitation in a Victor2 plate reader (PerkinElmer, Shelton, USA) at 30°C to avoid condensation, during two and a half hours. After reaching mid-exponential growth (OD_600_ 0.4), cells were treated with the indicated effector or the oxidative stress-inducing agent (H_2_O_2_) at the indicated concentrations and the incubation at 30°C resumed to 24 h. Bacterial growth was recorded as the optical density at 600 nm measured every 15 min. Data corresponds to two independent experiments with eight technical replicates each.

## ACKNOWLEDGEMENTS

We are indebted to Rafael Silva-Rocha (Ribeirão Preto Medical School – USP, Brazil) and Juhyun Kim (Imperial College London, UK) for constructing the LacIq-Ptrc and XylS/Pm expression cargoes. This work was funded by SETH (RTI2018-095584-B-C42) (MINECO/FEDER) and SyCoLiM (ERA-COBIOTECH 2018 - PCI2019-111859-2) Projects of the Spanish Ministry of Science and Innovation, the MADONNA (H2020-FET-OPEN-RIA-2017-1-766975), BioRoboost (H2020-NMBP-BIO-CSA-2018-820699), SynBio4Flav (H2020-NMBP-TR-IND/H2020-NMBP-BIO-2018-814650) and MIX-UP (MIX-UP H2020-BIO-CN-2019-870294) Contracts of the European Union and the InGEMICS-CM (S2017/BMD-3691) Project of the Comunidad de Madrid - European Structural and Investment Funds - (FSE, FECER) to V.D.L. and the Novo Nordisk Foundation (individual grant NNF10CC1016517, and LiFe, NNF18OC0034818), the European Union’s Horizon 2020 Research and Innovation Programme under grant agreement No. 814418 (SinFonia) and the Danish Council for Independent Research (SWEET, DFF-Research Project 8021-00039B) to P.I.N. Authors declare no conflict of interest.

## Supplementary Materials

**Supplementary Table S1** Strains and plasmids used in this study.

**Supplementary Table S2** Primers used in the PCR reactions

## REFERENCES

Amann, E., Ochs, B., and Abel, K.-J. (1988) Tightly regulated tac promoter vectors useful for the expression of unfused and fused proteins in *Escherichia coli*. Gene 69: 301–315.

Andrews, B., Adari, H., Hannig, G., Lahue, E., Gosselin, M., Martin, S. et al. (1996) A tightly regulated high level expression vector that utilizes a thermosensitive lac repressor: production of the human T cell receptor Vβ5.3 in *Escherichia coli*. Gene 182: 101–109.

Arce-Rodriguez, A., Benedetti, I., Silva-Rocha, R., and de Lorenzo, V. (2020) Standardization of inducer-activated broad host range expression modules: Debugging and refactoring an alkane-responsive AlkS/PalkB device. bioRxiv: 2020.2012.2026.424440.

Balzer, S., Kucharova, V., Megerle, J., Lale, R., Brautaset, T., and Valla, S. (2013) A comparative analysis of the properties of regulated promoter systems commonly used for recombinant gene expression in *Escherichia coli*. Microb Cell Fact 12: 26.

Baneyx, F. (1999) Recombinant protein expression in *Escherichia coli*. Curr Opinin Biotechnol 10: 411–421.

Bar-Even, A., Paulsson, J., Maheshri, N., Carmi, M., O’Shea, E., Pilpel, Y., and Barkai, N. (2006) Noise in protein expression scales with natural protein abundance. Nat Genet 38: 636–643.

Beal, J., Goñi-Moreno, A., Myers, C., Hecht, A., de Vicente, M.D.C., Parco, M. et al. (2020) The long journey towards standards for engineering biosystems: Are the Molecular Biology and the Biotech communities ready to standardise? EMBO Rep 21: e50521.

Benedetti, I. (2014) Design of standardized molecular tools to analyze regulatory properties and biotechnologial applications of the soli bacterium *Pseudomonas putida*. In Molecular Biology Department: Universidad Autónoma de Madrid.

Benedetti, I., Nikel, P.I., and de Lorenzo, V. (2016) Data on the standardization of a cyclohexanone-responsive expression system for Gram-negative bacteria. Data Brief 6: 738–744.

Bertram, R., and Hillen, W. (2008) The application of Tet repressor in prokaryotic gene regulation and expression. Microb Biotechnol 1: 2–16.

Blatny, J.M., Brautaset, T., Winther-Larsen, H.C., Haugan, K., and Valla, S. (1997) Construction and use of a versatile set of broad-host-range cloning and expression vectors based on the RK2 replicon. Appl Environ Microbiol 63: 370–379.

Brosius, J., Erfle, M., and Storella, J. (1985) Spacing of the −10 and −35 regions in the tac promoter. Effect on its in vivo activity. J Biol Chem 260: 3539–3541.

Cabiscol, E., Tamarit, J., and Ros, J. (2000) Oxidative stress in bacteria and protein damage by reactive oxygen species. Int Microbiol 3: 3–8.

Calero, P., Jensen, S.I., and Nielsen, A.T. (2016) Broad-Host-Range ProUSER Vectors Enable Fast Characterization of Inducible Promoters and Optimization of p-Coumaric Acid Production in *Pseudomonas putida* KT2440. ACS Synth Biol 5: 741–753.

Calos, M.P. (1978) DNA sequence for a low-level promoter of the lac repressor gene and an ‘up’ promoter mutation. Nature 274: 762–765.

Calles, B., Goñi-Moreno, Á., and de Lorenzo, V. (2019) Digitalizing heterologous gene expression in Gram-negative bacteria with a portable ON/OFF module. Mol Syst Biol 15: e8777.

Canton, B., Labno, A., and Endy, D. (2008) Refinement and standardization of synthetic biological parts and devices. Nat Biotechnol 26: 787–793.

Chalancon, G., Ravarani, C.N.J., Balaji, S., Martinez-Arias, A., Aravind, L., Jothi, R., and Babu, M.M. (2012) Interplay between gene expression noise and regulatory network architecture. Trends Genet 28: 221–232.

Choudhary, S., and Schmidt-Dannert, C. (2010) Applications of quorum sensing in biotechnology. Appl Microb Biotechnol 86: 1267–1279.

Egan, S.M., and Schleif, R.F. (1993) A Regulatory Cascade in the Induction of rhaBAD. J Mol Biol 234: 87–98.

Elowitz, M.B., Levine, A.J., Siggia, E.D., and Swain, P.S. (2002) Stochastic Gene Expression in a Single Cell. Science 297: 1183–1186.

Elvin, C.M., Thompson, P.R., Argall, M.E., Philip Hendr, N., Stamford, P.J., Lilley, P.E., and Dixon, N.E. (1990) Modified bacteriophage lambda promoter vectors for overproduction of proteins in *Escherichia coli*. Gene 87: 123–126.

Ellefson, J.W., Ledbetter, M.P., and Ellington, A.D. (2018) Directed evolution of a synthetic phylogeny of programmable Trp repressors. Nat Chem Biol 14: 361–367.

Gawin, A., Valla, S., and Brautaset, T. (2017) The XylS/Pm regulator/promoter system and its use in fundamental studies of bacterial gene expression, recombinant protein production and metabolic engineering. Microb Biotechnol 10: 702–718.

Glascock, C.B., and J. Weickert, M. (1998) Using chromosomal lacIQ1 to control expression of genes on high-copy-number plasmids in *Escherichia coli*. Gene 223: 221–231.

Goñi-Moreno, Á., Benedetti, I., Kim, J., and de Lorenzo, V. (2017) Deconvolution of Gene Expression Noise into Spatial Dynamics of Transcription Factor–Promoter Interplay. ACS Synth Biol 6: 1359–1369.

Guzman, L.M., Belin, D., Carson, M.J., and Beckwith, J. (1995) Tight regulation, modulation, and high-level expression by vectors containing the arabinose PBAD promoter. J Bacteriol 177: 4121–4130.

Haldimann, A., Daniels, L.L., and Wanner, B.L. (1998) Use of New Methods for Construction of Tightly Regulated Arabinose and Rhamnose Promoter Fusions in Studies of the *Escherichia coli* Phosphate Regulon. J Bacteriol 180: 1277–1286.

Henßler, E.-M., Bertram, R., Wisshak, S., and Hillen, W. (2005) Tet repressor mutants with altered effector binding and allostery. FEBS J 272: 4487–4496.

Hernández-Arranz, S., Sánchez-Hevia, D., Rojo, F., and Moreno, R. (2016) Effect of Crc and Hfq proteins on the transcription, processing, and stability of the *Pseudomonas putida* CrcZ sRNA. Rna 22: 1902–1917.

Heyneker, H.L., Shine, J., Goodman, H.M., Boyer, H.W., Rosenberg, J., Dickerson, R.E. et al. (1976) Synthetic lacoperator DNA is functional *in vivo*. Nature 263: 748–752.

Hilfinger, A., and Paulsson, J. (2011) Separating intrinsic from extrinsic fluctuations in dynamic biological systems. Proc Nat Acad Sci 108: 12167–12172.

Inouye, S., Nakazawa, A., and Nakazawa, T. (1987a) Expression of the regulatory gene xylS on the TOL plasmid is positively controlled by the xylR gene product. Proc Natl Acad Sci 84: 5182–5186.

Inouye, S., Nakazawa, A., and Nakazawa, T. (1987b) Overproduction of the xylS gene product and activation of the xylDLEGF operon on the TOL plasmid. J Bacteriol 169: 3587–3592.

Kærn, M., Elston, T.C., Blake, W.J., and Collins, J.J. (2005) Stochasticity in gene expression: from theories to phenotypes. Nat Rev Genet 6: 451–464.

Kaur, J., Kumar, A., and Kaur, J. (2018) Strategies for optimization of heterologous protein expression in *E. coli*: Roadblocks and reinforcements. Int J Biol Macromol 106: 803–822.

Kelly, C.L., Liu, Z., Yoshihara, A., Jenkinson, S.F., Wormald, M.R., Otero, J. et al. (2016) Synthetic Chemical Inducers and Genetic Decoupling Enable Orthogonal Control of the rhaBAD Promoter. ACS Synth Biol 5: 1136–1145.

Kemp, L.R., Dunstan, M.S., Fisher, K., Warwicker, J., and Leys, D. (2013) The transcriptional regulator CprK detects chlorination by combining direct and indirect readout mechanisms. Phill Trans R Soc B 368: 20120323.

Kent, R., and Dixon, N. (2020) Contemporary Tools for Regulating Gene Expression in Bacteria. Trends Biotech 38: 316–333.

Kessler, B., Herrero, M., Timmis, K.N., and de Lorenzo, V. (1994) Genetic evidence that the XylS regulator of the *Pseudomonas* TOL meta operon controls the Pm promoter through weak DNA-protein interactions. J Bacteriol 176: 3171–3176.

Korpela, M.T., Kurittu, J.S., Karvinen, J.T., and Karp, M.T. (1998) A Recombinant *Escherichia coli* Sensor Strain for the Detection of Tetracyclines. Anal Chem 70: 4457–4462.

Lanzer, M., and Bujard, H. (1988) Promoters largely determine the efficiency of repressor action. Proc Natl Acad Sci 85: 8973–8977.

Lestas, I., Paulsson, J., Ross, N.E., and Vinnicombe, G. (2008) Noise in Gene Regulatory Networks. IEEE Trans Autom Control 53: 189–200.

Love, C.A., Lilley, P.E., and Dixon, N.E. (1996) Stable high-copy-number bacteriophage λ promoter vectors for overproduction of proteins in *Escherichia coli*. Gene 176: 49–53.

Marqués, S., and Ramos, J.L. (1993) Transcriptional control of the *Pseudomonas putida* TOL plasmid catabolic pathways. Mol Microbiol 9: 923–929.

Marschall, L. (2017) Tunable recombinant protein expression in *E. coli*: promoter systems and genetic constraints. App microb biotechnol 101: pp. 501–512-2017 v.2101 no.2012.

Mermod, N., Ramos, J.L., Bairoch, A., and Timmis, K.N. (1987) The xylS gene positive regulator of TOL plasmid pWWO: Identification, sequence analysis and overproduction leading to constitutive expression of meta cleavage operon. Mol Gen Genet MGG 207: 349–354.

Meyer, A.J., Segall-Shapiro, T.H., Glassey, E., Zhang, J., and Voigt, C.A. (2019) *Escherichia coli* “Marionette” strains with 12 highly optimized small-molecule sensors. Nat Chem Biol 15: 196–204.

Michan, C., Zhou, L., Gallegos, M.T., Timmis, K.N., and Ramos, J.L. (1992) Identification of critical amino-terminal regions of XylS. The positive regulator encoded by the TOL plasmid. J Biol Chem 267: 22897–22901.

Miller, W.G., and Lindow, S.E. (1997) An improved GFP cloning cassette designed for prokaryotic transcriptional fusions. Gene 191: 149–153.

Munsky, B., Neuert, G., and van Oudenaarden, A. (2012) Using Gene Expression Noise to Understand Gene Regulation. Science 336: 183–187.

Nikel, P.I., and de Lorenzo, V. (2018) *Pseudomonas putida* as a functional chassis for industrial biocatalysis: From native biochemistry to trans-metabolism. Metab Eng 50: 142–155.

Oehler, S., Eismann, E.R., Krämer, H., and Müller-Hill, B. (1990) The three operators of the lac operon cooperate in repression. EMBO J 9: 973–979.

Oehler, S., Amouyal, M., Kolkhof, P., von Wilcken-Bergmann, B., and Müller-Hill, B. (1994) Quality and position of the three lac operators of *E. coli* define efficiency of repression. EMBO J 13: 3348–3355.

Paulsson, J. (2004) Summing up the noise in gene networks. Nature 427: 415–418.

Pedraza, J.M., and van Oudenaarden, A. (2005) Noise Propagation in Gene Networks. Science 307: 1965–1969.

Ramos, J.L., Mermod, N., and Timmis, K.N. (1987) Regulatory circuits controlling transcription of TOL plasmid operon encoding meta-cleavage pathway for degradation of alkylbenzoates by *Pseudomonas*. Mol Microbiol 1: 293–300.

Ramos, J.L., Michan, C., Rojo, F., Dwyer, D., and Timmis, K. (1990) Signal-regulator interactions, genetic analysis of the effector binding site of xyls, the benzoate-activated positive regulator of *Pseudomonas* TOL plasmid meta-cleavage pathway operon. J Mol Biol 211: 373–382.

Ramos, J.L., Martínez-Bueno, M., Molina-Henares, A.J., Terán, W., Watanabe, K., Zhang, X. et al. (2005) The TetR Family of Transcriptional Repressors. Microbiol Mol Biol Rev 69: 326–356.

Rojo, F. (2009) Degradation of alkanes by bacteria. Environ Microbiol 11: 2477–2490.

Sambrook, J., Fritsch, E. F., and Maniatis, T. (1989) Molecular cloning: A Laboratory Manual: Cold Spring Harbor, NY.

Sanchez, A., Choubey, S., and Kondev, J. (2013) Regulation of Noise in Gene Expression. Annu Rev Biophys 42: 469–491.

Scholz, O., Köstner, M., Reich, M., Gastiger, S., and Hillen, W. (2003) Teaching TetR to Recognize a New Inducer. J Mol Biol 329: 217–227.

Schuller, A., Cserjan-Puschmann, M., Tauer, C., Jarmer, J., Wagenknecht, M., Reinisch, D. et al. (2020) *Escherichia coli* σ70 promoters allow expression rate control at the cellular level in genome-integrated expression systems. Microb Cell Fact 19: 58.

Silva-Rocha, R., Martínez-García, E., Calles, B., Chavarría, M., Arce-Rodríguez, A., de las Heras, A. et al. (2012) The Standard European Vector Architecture (SEVA): a coherent platform for the analysis and deployment of complex prokaryotic phenotypes. Nucleic Acids Res 41: D666–D675..

Skerra, A. (1994) Use of the tetracycline promoter for the tightly regulated production of a murine antibody fragment in *Escherichia coli*. Gene 151: 131–135.

Sørensen, H.P., and Mortensen, K.K. (2005) Advanced genetic strategies for recombinant protein expression in *Escherichia coli*. J Biotechnol 115: 113–128.

Spooner, R.A., Bagdasarian, M., and Franklin, F.C. (1987) Activation of the xylDLEGF promoter of the TOL toluene-xylene degradation pathway by overproduction of the xylS regulatory gene product. J Bacteriol 169: 3581–3586.

Steigedal, M., and Valla, S. (2008) The Acinetobacter sp. chnB promoter together with its cognate positive regulator ChnR is an attractive new candidate for metabolic engineering applications in bacteria. Metab Eng 10: 121–129.

Studier, F.W., and Moffatt, B.A. (1986) Use of bacteriophage T7 RNA polymerase to direct selective high-level expression of cloned genes. J Mol Biol 189: 113–130.

Swain, P.S., Elowitz, M.B., and Siggia, E.D. (2002) Intrinsic and extrinsic contributions to stochasticity in gene expression. Proc Nat Acad Sci 99: 12795–12800.

Tang, S.-Y., and Cirino, P.C. (2011) Design and Application of a Mevalonate-Responsive Regulatory Protein. Angew Chem Int Ed 50: 1084–1086.

Tang, S.-Y., Fazelinia, H., and Cirino, P.C. (2008) AraC Regulatory Protein Mutants with Altered Effector Specificity. J Am Chem Soc 130: 5267–5271.

Tas, H., Grozinger, L., Stoof, R., de Lorenzo, V., and Goñi-Moreno, Á. (2021) Contextual dependencies expand the re-usability of genetic inverters. Nat Commun 12: 355.

Taylor, N.D., Garruss, A.S., Moretti, R., Chan, S., Arbing, M.A., Cascio, D. et al. (2016) Engineering an allosteric transcription factor to respond to new ligands. Nat Methods 13: 177–183.

Terpe, K. (2006) Overview of bacterial expression systems for heterologous protein production: from molecular and biochemical fundamentals to commercial systems. App Microbiol Biotechnol 72: 211–222.

van Beilen, J.B., Panke, S., Lucchini, S., Franchini, A.G., Röthlisberger, M., and Witholt, B. (2001) Analysis of *Pseudomonas putida* alkane-degradation gene clusters and flanking insertion sequences: evolution and regulation of the alk genes. Microbiology 147: 1621–1630.

Wu, H.-Y., Shyy, S., Wang, J.C., and Liu, L.F. (1988) Transcription generates positively and negatively supercoiled domains in the template. Cell 53: 433–440.

Yin, J., Li, G., Ren, X., and Herrler, G. (2007) Select what you need: A comparative evaluation of the advantages and limitations of frequently used expression systems for foreign genes. J Biotechnol 127: 335–347.

Zerbs, S., Frank, A.M., and Collart, F.R. (2009) Chapter 12 Bacterial Systems for Production of Heterologous Proteins. In Methods in Enzymology. Burgess, R.R., and Deutscher, M.P. (eds): Academic Press, pp. 149–168.

Zhao, X., and Drlica, K. (2014) Reactive oxygen species and the bacterial response to lethal stress. Curr Opin Microbiol 21: 1–6.

