## Supplementary material for "Parameterization of regulatory nodes for engineering broad host range heterologous gene expression": Supllemental Table S1 and S2

**Supplementary Table S1.** Strains and plasmids used in this study

| Strain or plasmid | Description | References and/or source |
|---|---|---|
| Strain (E. coli) | | |
| DH10B | $F^-$ *endA1 recA1 galE15 galK16 nupG rpsL ΔlacX74 Φ80lacZΔM15 araD139 Δ(ara,leu)7697 mcrA Δ(mrr-hsdRMS-mcrBC)* $λ^-$ | Invitrogen |
| DH5α | *supE44, ΔlacU169, (ϕ80 lacZDM15), hsdR17, (rk$^-$ mk$^+$), recA1, endA1, thi1, gyrA, relA* | Invitrogen |
| CC118 | $F^-$, Δ*(ara-leu)7697, araD139, Δ(lac)X74, phoAΔ20, galE, galK, thi, rpsE, rpoB, argE(Am), recA1* | (Manoil and Beckwith, 1985) |
| Plasmid | Relevant features* | |
| pGA-LacI | Derivative of plasmid p57 inserted with a synthetic fragment containing edited msf•GFP gene. $Ap^R$; *ori* MB1 | GeneArt® |
| pS237M | $Km^R$; *oriV* (pBBR1), promoterless msf•GFP | (Calles et al., 2019; Benedetti et al., 2016) |
| pTRrc-99A | trcP vector, $lacI^q$, pUC18 EcoRI-HindIII polylinker region | (Amann et al., 1988) |
| pS224 | Derivative of pSEVA221 bearing a *lacI$^q$*-$P_{Trc}$ expression cassette, $Km^R$; ori RK2, standard multiple cloning site | (Silva-Rocha et al., 2012) |
| pSEVA234 | Derivative of pSEVA221 bearing a *lacI$^q$* -PTrc expression cassette, $Km^R$; *ori* pBBR1, standard multiple cloning site | This work |
| pS234M | pSEVA 234 derivative, $Km^R$; *oriV* (pBBR1), IPTG inducible msf•GFP | This work |
| pJB653 | RK2 expression vector containing the *Pm* promoter and the gene encoding XylS. *oriV*, $Ap^R$, 7 Kb | (Blatny et al., 1997) |

| | | |
|---|---|---|
| pSEVA226 | Derivative of pSEVA221 bearing a promoterless *luxCDABE* operon, $Km^R$; ori RK2, standard multiple cloning site | (Silva-Rocha et al., 2012) |
| pSEVA226-XylS-Pm | Derivative of pSEVA226. $Km^R$; oriV (pBBR1); cargo (*xylS-Pm* → *luxCDABE*) | (Silva-Rocha et al., 2012) |
| pSEVA238 | Inducible expression vector; $Km^R$, oriV (pBBR1); *xylS/Pm*; standard multiple cloning site | (Calles et al., 2019) |
| pS238M | pSEVA238 derivative bearing the msf•GFP gene; $Km^R$; *oriV* (pBBR1); cargo (*xylS-Pm* → *msf•GFP*) | (Calles et al., 2019) |
| pSEVA429 | Inducible expression vector; $Sm^R$; *oriV* (RK2); *alkS/*$P_{alkB}$; standard multiple cloning site | (Arce-Rodriguez et al., 2020) |
| pSEVA239 | Inducible expression vector; $Km^R$, *oriV* (pBBR1); *alkS/*$P_{alkB;}$ standard multiple cloning site | This work |
| pS239M | pSEVA239 derivative bearing the msf•GFP gene; $Km^R$; *oriV* (PBBR1); cargo (*alkS/PalkB* → *msf•GFP*) | (Calles et al., 2019) |
| pSEVA2311 | *oriV* (pBBR1), *chnR,* $P_{chnB}$; standardized, cyclohexanone-responsive expression vector | (Benedetti et al., 2016) |
| pS2311M ( = pSCM) | $Km^R$; *oriV* (pBBR1), *chnR,* cargo (*PchnB*→ *msf•GFP*) | (Benedetti et al., 2016) |
| pMLK1 | $Ap^R$, *oriV* ColE1, pET28a carrying CpRK200S (*cprK1*) of *Desulfitobacterium hafniense* | (Kemp et al., 2013) |
| pSEVA2312 | Inducible expression vector; $Km^R$, *oriV* (pBBR1); *cprK/*$P_{DB3};$ standard multiple cloning site | (Benedetti, 2014) |
| pS2312M | $Km^R$; *oriV* (pBBR1), *cprK1* cargo (*PchnB*→ *msf•GFP*) | This work |

*The abbreviations used in this table are as follows: Km, kanamycin; Ap, ampicillin; msf•GFP, gene encoding the monomeric and superfolder green fluorescent protein. Plasmids from the SEVA collection are freely available in the SEVA database (http://seva-plasmids.com/)

**Supplementary Table S2.** Primers used in the PCR reactions

| Oligonucleotide | Sequence* |
|---|---|
| TRC-F | CCGGCCTAGGGTGTGAAATTGTTATCCGCTCAC |
| TRC-R | GGCGTTAATTAATTGACACCATCGAATGGTGC |
| msfGFPHindIII-F | TATATAAGCTTAGGAGGAAAAACATATGCGTAAAGGTGAAGAACTGTTC |
| msfGFPSpeI-R | GATATACTAGTTTATTTGTAGAGTTCATCCATG |
| 5-CPRK1 | GCGCGCCTAGGGCTAGCATGGCTGTTGAAGGTTTGGGCAAG |
| 3-CPRK1 | CGCGCTTAATTAACTAGTAATACGATGTTTGTTCAG |

* Sequences of oligonucleotides employed in this study. Restriction sites entered for cloning purposes are underlined.

REFERENCES

Amann, E., Ochs, B., and Abel, K.-J. (1988) Tightly regulated tac promoter vectors useful for the expression of unfused and fused proteins in *Escherichia coli*. *Gene* **69**: 301-315.

Arce-Rodriguez, A., Benedetti, I., Silva-Rocha, R., and de Lorenzo, V. (2020) Standardization of inducer-activated broad host range expression modules: Debugging and refactoring an alkane-responsive AlkS/PalkB device. *bioRxiv*: 2020.2012.2026.424440.

Benedetti, I. (2014) Design of standardized molecular tools to analyze regulatory properties and biotechnologial applications of the soli bacterium *Pseudomonas putida*. In *Molecular Biology Department*: Universidad Autónoma de Madrid.

Benedetti, I., Nikel, P.I., and de Lorenzo, V. (2016) Data on the standardization of a cyclohexanone-responsive expression system for Gram-negative bacteria. *Data Brief* **6**: 738-744.

Blatny, J.M., Brautaset, T., Winther-Larsen, H.C., Haugan, K., and Valla, S. (1997) Construction and use of a versatile set of broad-host-range cloning and expression vectors based on the RK2 replicon. *Appl Environ Microbiol* **63**: 370-379.

Calles, B., Goñi-Moreno, Á., and de Lorenzo, V. (2019) Digitalizing heterologous gene expression in Gram-negative bacteria with a portable ON/OFF module. *Mol Syst Biol* **15**: e8777.

Kemp, L.R., Dunstan, M.S., Fisher, K., Warwicker, J., and Leys, D. (2013) The transcriptional regulator CprK detects chlorination by combining direct and indirect readout mechanisms. *Phill Trans R Soc B* **368**: 20120323.

Manoil, C., and Beckwith, J. (1985) TnphoA: a transposon probe for protein export signals. *Proc Nat Acad Sci* **82**: 8129-8133.

Silva-Rocha, R., Martínez-García, E., Calles, B., Chavarría, M., Arce-Rodríguez, A., de las Heras, A. et al. (2012) The Standard European Vector Architecture (SEVA): a coherent platform for the analysis and deployment of complex prokaryotic phenotypes. *Nucleic Acids Res* **41**: D666-D675..
